# Splicing conservation signals in plant long non-coding RNAs

**DOI:** 10.1101/588954

**Authors:** Jose Antonio Corona-Gomez, Irving Jair Garcia-Lopez, Peter F. Stadler, Selene L. Fernandez-Valverde

## Abstract

Long non-coding RNAs (lncRNAs) have recently emerged as prominent regulators of gene expression in eukaryotes. LncRNAs often drive the modification and maintenance of gene activation or gene silencing states via chromatin conformation rearrangements. In plants, lncRNAs have been shown to participate in gene regulation, and are essential to processes such as vernalization and photomorphogenesis. Despite their prominent functions only over a dozen lncRNAs have been experimentally and functionally characterized.

Similar to its animal counterparts, the rates of sequence divergence are much higher in plant lncRNAs than in protein coding mRNAs, making it difficult to identify lncRNA conservation using traditional sequence comparison methods. Beyond this, little is known about the evolutionary patterns of lncRNAs in plants. Here, we characterized the splicing conservation of lncRNAs in Brassicaceae. We generated a whole-genome alignment of 16 Brassica species and used it to identify synthenic lncRNA orthologues. Using a scoring system trained on transcriptomes from *A. thaliana* and *B. oleracea*, we identified splice sites across the whole alignment and measured their conservation. Our analysis revealed that 17.9% (112/627) of all intergenic lncRNAs display splicing conservation in at least one exon, an estimate that is substantially higher to previous estimates of lncRNA conservation in this group. Our findings agree with similar studies in vertebrates, demonstrating that splicing conservation can be evidence of stabilizing selection. We provide conclusive evidence for the existence of evolutionary deeply conserved lncRNAs in plants and describe a generally applicable computational workflow to identify functional lncRNAs in plants.

## 1. INTRODUCTION

Long non-coding RNAs (lncRNAs), by definition, do not code for proteins. Over the last decade, a wide variety of mechanisms have been discovered by which lncRNAs contribute to the regulation of the expression of protein-coding genes and small RNAs (Liu et al. (2015); Chekanova (2015); Ulitsky (2016); Wang Chekanova (2017); Yamada (2017)). Most lncRNAs are found in the nucleus associated with the chromatin, regulating gene expression by recruiting components of the epigenetic machinery to specific genomic locations. Some lncRNAs also influence genome stability and nuclear domain organization. Serving as molecular sponges and decoys, they act both at the transcriptional level, by affecting RNA-directed DNA methylation; in post-transcriptional regulation, by inhibiting the interaction between microRNAs (miRNAs) and their target messenger RNAs (mRNAs); and by controlling alternative splicing due to sequestration of splicing factors (Bardou et al., 2014). Hence they differ not only in size but also in their biogenesis and molecular mechanisms from small RNAs such as miRNAs and siRNAs (Bánfai et al., 2012). lncRNAs are regulated and processed similar to mRNAs (Mercer Mattick, 2013) and their expression patterns are often very specific to particular tissues or developmental stages (Mercer Mattick, 2013). Recent data suggest that there appears to be a distinction between highly conserved, constitutively transcribed lncRNAs and tissue-specific lncRNAs with low expression levels (Deng et al., 2018b; Sarropoulos et al., 2019).

Despite their often very poor sequence conservation (Necsulea et al., 2014), the majority of lncRNAs are well-conserved across animals, as evidenced by the conservation of many of their splice sites (Nitsche et al., 2015). While well-conserved as entities, they show much more plasticity in their gene structure and sequence than protein-coding genes. The many lineagespecific differences have implicated lncRNAs as major players in lineage-specific adaptation (Lozada-Chávez et al., 2011): changes in transcript structure are likely associated with the inclusion or exclusion of sets of protein or miRNA binding sites and hence may have large effects on function and specificity of a particular lncRNA.

The systematic annotation of orthologous lncRNAs is important not only to provide reasonably complete maps of the transcriptome, but also as means of establishing that a particular lncRNA has a biological function. After all, conservation over long evolutionary timescales is often used as the most important argument for the biological function of an open reading frame in the absence of direct experimental evidence for translation and experimental data characterizing the peptide product. While a large amount of work is available showing that vertebrate genomes contain a large number of secondary elements that are under negative selection (Seemann et al., 2017; Smith et al., 2013; Hezroni et al., 2015; Nitsche Stadler, 2017) and the majority of human lncRNAs are evolutionary old (Nitsche et al., 2015), a much less systematic and complete picture is available for plants.

In fact, detailed studies into the evolution of plant lncRNAs have been rare until very recently. An analysis of lncRNAs in five monocot and five dicot species (Deng et al., 2018b) found that the majority of lncRNAs are poorly conserved at sequence level while a majority is highly divergent but syntenically conserved. These positionally conserved lncRNAs were previously found to be located near telomeres in *A. thaliana* (Mohammadin et al., 2015). Plant lncRNAs have also been shown to display canonical splicing signals (Deng et al., 2018b). Another study in 10 Brassicaceae genomes found 22% conservation of intergenic lncRNA loci (Nelson et al., 2016), as well as little evidence of an impact of whole genome duplications or transposable element (TE) activity on the emergence of lincRNAs.

Nevertheless, there are some plant lncRNAs whose regulatory functions have been studied extensively and are understood at a level of detail comparable to most proteins (Rai et al., 2019): *COOLAIR* in Brassicaceae has a crucial role in the vernalization process (Hawkes et al., 2016) and its transcription accelerates epigenetic silencing of the flowering locus C (FLC) (Rosa et al., 2016). The lncRNA *HID1* is a key component in promoting photomorphogenesis in response to different levels of red light (Wang et al., 2014). *HID1* is highly conserved and acts through binding to chromatin in *trans* to act upon the *PIF3* promoter. A similar trans-acting lncRNA is *ELENA1*, which functions in plant immunity (Mach, 2017). Competing endogenous RNAs (ceRNAs) acts as “sponges” for miRNAs. In plants, ceRNAs are a large class of lncRNAs (Yuan et al., 2017; Paschoal et al., 2018) and form extensive regulatory networks (Meng et al., 2018; Zhang et al., 2018). The paradigmatic example in *A. thaliana* is *IPS1*, which sequesters miR399, resulting in changes in phosphate homeostasis (Franco-Zorrilla et al., 2007).

Although the functional characterization of plant lncRNAs is confined to a small number of cases, plant lncRNAs are being reported at a rapidly increasing pace (Nelson et al., 2016). As in the case of animals, it is important therefore to amass evidence for the functionality of individual transcripts. Differential expression, or correlations with important regulatory proteins or pathways alone do not provide sufficient evidence to decide whether a transcript has a causal effect or whether its expression pattern is a coincidental downstream effect. As a first step towards prioritizing candidates for functional characterization, we advocate for the use of unexpectedly deep conservation of the gene structure as an indicator of biological function. While logically this still does not inform about function in an specific context, it is much less likely that changes in expression patterns of a conserved and thus presumably functional molecule are without biological consequence.

The much higher level of plasticity in plant genomes, compared to animal genomes, potentially makes it more difficult to trace the evolution of lncRNAs. We therefore concentrate here on a phylogenetically relatively narrow group, the Brassicaceae, with genomes that are largely alignable with each other. We track the conservation of functional elements, in particular splice junctions, through the entire data set. This provides direct evidence also in cases where transcriptome data is not available in sufficient coverage and or sufficient diversity of tissues and/or developmental stages. As a final result, we provide a list of homologous lncRNAs in Brassicaceae as well as a detailed map of the conservation of splice sites in this clade.

## 2. MATERIALS AND METHODS

### 2.1. Whole genome alignment

We selected sixteen plant genomes from those available for the Brassicaceae family in NCBI, Phytozome, and Ensembl-Plants (Suppl. Tab. S1) based on the quality of assembly, as measured by the number of contigs/scaffolds. All genomes were downloaded in fasta format. Mitochondrial and chloroplast sequences were excluded based on annotation.

The genomes were aligned using `Cactus` v0 (Paten et al., 2011). Like other whole genome alignments (WGA) methods, `Cactus` v0 uses small regions with very high sequence similarity as anchors. To resolve conflicts at this level, `Cactus` v0 uses a specialized graph data structure that produces better overall alignments than other WGA approaches (Earl et al., 2014). The final WGA result were stored in HAL format (Hickey et al., 2013) for further processing.

### 2.2. Transcriptome data and assembly

We used four previously published base-line transcriptomes for *A. thaliana* (Liu et al., 2012) (GEO accession number GSE38612), as well as transcriptomes of shade response experiments from (Kohnen et al., 2016) (GEO accession number GSE81202). For *Brassica oleracea* we used transcriptomes from (Yu et al., 2014) (Expression Atlas accession number E-GEOD-42891). To validate predicted lncRNAs we used the publicly available transcriptome data sets listed in Suppl. Tab. S2. All transcriptomes were downloaded as raw reads in `fastq` format.

We generated our own lncRNA annotation using all single-end stranded sequencing libraries from (Kohnen et al., 2016). Libraries were quality-filtered using `Trimmomatic` v0.32 (Bolger et al., 2014), and mapped to the TAIR10 genome (Berardini et al., 2015) using `tophat v2.1.1` (Trapnell et al., 2009) with parameters: `-I 20 -I 1000 -read-edit-dist 3` `-read-realign-edit-dist 0 -library-type` `fr-firstsrand -g 1`. Transcripts were assembled with `Cufflinks v2.2.1` (Trapnell et al., 2010) with parameters: `--overlap-radius 1 -p 8 -I 1000` `-min-intron-length 20 -g TAIR10 GFF3.gff` `-library-type fr-firststrand` and subsequently merged into a single reference transcriptome using `Cuffmerge` v2.2.1.

### 2.3. lncRNA Annotation

LncRNAs in the (Kohnen et al., 2016) dataset were annotated using two independent methods. First, coding and non-coding transcripts were identified with `CPC` v0.9.r2 (Coding Potential Calculator) (Kong et al., 2007), a support vector machine classifier. Additionally, we used a strict stepwise annotation workflow (Cabili et al., 2011) on all transcripts. Specifically, we removed transcripts less than 200 nt in length, and identified ORFs 75 aminoacids or longer. Identified ORFs were compared against the NCBI non redundant (nr) database using `blastx` v2.2.31 and `blastp` v2.2.31 (Altschul et al., 1990) with *E-value* and cutoff of *E* < 10 for a sequence to be considered potentially coding. In addition, we used `HMMER` v3.1b2 (Wheeler Eddy, 2013) to search for `Pfam` protein domains, `signalP` v4.1 (Petersen et al., 2011) to identify signal peptides, and `tmhmm` v2.0 (Krogh et al., 2001) for transmembrane helices. Only sequences that had no similarity with proteins in nr and no identifiable protein domains, signal peptides or transmembrane domains were annotated as *bona fide* lncRNAs.

To characterize the genomic context of identified lncRNAs, we used `bedtools` v2.25.0 (Quinlan Hall, 2010) and compared the lncRNA annotation with the protein coding gene annotation in Araport11 (Cheng et al., 2017). All lncRNA candidates that overlapped a coding sequence or some other ncRNA (miRNA, snoRNA, snRNA) by at least 1 nt were discarded.

### 2.4. Splicing map

The construction of splicing maps requires a seed set of experimentally determined splice sites in at least one species as well as a statistical model to assess the conservation of splice donors and splice acceptors whenever no direct experimental evidence is available.

To obtain these data for Brassicaceae, we mapped the reference transcriptomes to the corresponding reference genome using `STAR` v2.4.0.1 (Dobin et al., 2013) with default parameters. The table of splice junctions produced by `STAR` v2.4.0.1 for each data set were concatenated. Only splice junctions that (a) had at least 10 uniquely mapped reads crossing the junction, and (b) showed the canonical `GT`/`AG` dinucleotides delimiting the intron (c) within an intron of size between 59 bp and 999 bp were retained for subsequent analyses. Since some of the transcriptome datasets were not strand-specific, we included `CT`/`AC` delimiters, interpreting these as reverse-complements. The same procedure was used for splice site validation in other species, where each transcriptome was mapped against their respective genomes prior to splice junction identification. See Supplemental Table S1 for accessions.

For each identified splice site in *A. thaliana*, we used the `HalTools` v2.1 liftover tool (Hickey et al., 2013) to determine the corresponding orthologous positions in all other genome sequences in the `Cactus` v0 generated WGA. For each of the retained splicesite we extracted the genomic sequences surrounding the donor and acceptor sites. If more than one homolog per species is contained in the WGA, we retained the candidate with the highest sequence similarity to *A. thaliana*. For each known splice site and their orthologous position, the `MaxEntScan` v0 splicesite score (MES) (Yeo Burge, 2004) was computed with either the donor or acceptor model provided the region contained neither gaps nor ambiguous nucleotides (Suppl. Fig. S1). Otherwise, the regions was treated as non-conserved. `MaxEntScan` v0 models sequence motifs with a probabilistic model based on the Maximum Entropy Principle, which considers adjacent and non-adjacent dependencies between positions. Several works have verified that the MES is an informative score to measure splice site conservation (Nitsche et al., 2015; Eng et al., 2004). A `MaxEntScan` v0 splice-site score cutoff of 0 was used (Suppl. Fig. S2). This cut-off value was estimated from the distribution of the MES values obtained from *A. thaliana* and *B. oleracea* transcriptome data (Suppl. Fig. S2). To estimate the rate of false positives we calculated the probability of finding random splice sites in coding genes and in lincRNAs. For this, we sampled 10,000 random splice positions for both acceptor and donor splicing motifs. In addition to this we calculate the MES values of the same random positions conserved in all WGA species, verifying that they follow the same distribution as in *A. thaliana*. All positively predicted splice-sites, *i.e*., those with *MES* > 0, were added to the splicing map. The pipeline implementing this analysis is available at: bitbucket.org/JoseAntonioCorona/splicing map plants.

### 2.5. Data Availability

TrackHubs for all datasets and lncRNAs used in this study as well as WGA are available here: www.bioinf.uni-leipzig.de/Publications/SUPPLEMENTS/19-001/BrassicaceaeWGA/hub.txt

Additional information and machine readable intermediate results are provided at http://www.bioinf.uni-leipzig.de/Publications/SUPPLEMENTS/19-001

## 3. RESULTS

### 3.1. Identification of splice sites and lncRNAs

To build a *A. thaliana* splice junction reference, we identified about 125,000 introns using the transcriptomes of (Liu et al., 2012) compared with 175,000 introns annotated in TAIR10 (Release 38) (Berardini et al., 2015). The smaller number was expected as (i) only introns with convincing coverage by uniquely mapping reads were considered and (ii) not all *A. thaliana* genes are expressed in these four transcriptomes. Consistent with previous reports (Hebsgaard, 1996; Brown et al., 1996), the vast majority of the detected splice junctions have the canonical `GT`/`AG` motif required for inclusion into our splice site map. In total, we identified 222,772 individual sites in *A. thaliana* (117,644 donor and 121,002 acceptor sites). 55% of all donors but only 13% of the acceptors have aligned sequences in the WGA (Supp. Fig. S3). In addition, many splice sites have evidence of expression in transcriptome data from other species (Suppl. Tab. S3.)

To characterize splicing conservation in lncRNAs, we focused solely on intergenic long non-coding RNAs (lincRNAs). Conservation of splice sites in lncRNAs overlapping with coding genes may be confounded by the coding gene conservation signal, resulting in false positives. The lncRNAs described by Liu et al. (2012) comprise 595 lincRNAs with predicted introns, with only 18 with confirmed introns as annotated in Araport9 (Liu et al., 2012), while in Araport11 (Cheng et al., 2017) 288 annotated lincRNAs out of 2,444 have introns. We also used an additional set of lncRNAs expressed in *A. thaliana* cotyledons and hypocotyls in Col-0 plants in normal light or shade conditions (Kohnen et al., 2016). These libraries were stranded, and had three replicates as well as sufficient depth to produce a high confidence lncRNA annotation. As these transcriptomes are only derived from two experimental conditions (shadow and light) (Kohnen et al., 2016), they encompass only a fraction of the lncRNAs expressed throughout the *A. thaliana* life cycle. We identified 2,375 lncRNA transcripts, 1,465 of which overlapped with protein coding RNAs, while 808 were found in intergenic regions and were thus considered *bona fide* lincRNAs. In our analysis, we found 159 lincRNAs that were included in neither Araport11 nor TAIR10 (Cheng et al., 2017; Berardini et al., 2015). Furthermore, we excluded all lncRNAs that had any overlap with other annotated ncRNAs thus depleting our set of lncRNAs that may be microRNA or snoRNA precursors, as small RNAs are generally conserved. All 808 lincRNAs transcripts aggregated in 627 lincRNA genes, of which 58 have multiple isoforms. In contrast to the situation in animals, lincRNAs are therefore mostly mono-exonic in *A. thaliana*. Of the 627 lincRNA genes, only 173 had at least one intron and thus were used to test splice site conservation in lncRNAs; of these 173, only 35 were previously annotated in the Araport11 database.

### 3.2. Conservation of lncRNAs

To identify conserved elements by position we extracted aligned sequences corresponding to different annotations sets from the WGA. Between 69.6% to 44.2% of the *A. thaliana* genome was aligned with other Brassicaceae species. For the protein-coding genes annotated in Araport11 (Cheng et al., 2017) the alignment recovery rate ranges from 95.3% (26,153/27,445) (*A. lyrata*) to 86.9% (23,856/27,445) (*Aethionema arabicum*). As expected, the values are substantially lower for the Araport11 lincRNAs, where we recover between 77.1% (1,885/2,444) in *A. lyrata* and 50.8% (1243/2444) in *A. arabicum*. Using our own annotation, we recover between 62.0% (389/627) in *A. lyrata* and 38.1% (239/627) in *A. arabicum*, i.e., values comparable to the overall coverage of the genome. This reflects the fact that lncRNA sequences experience very little constraint on their sequence. Conservation (as measured by alignability) is summarized in Fig. 1 for different types of RNA elements. These values are comparable to a previous estimate of about 22% of the lincRNA loci are at least partially conserved at the sequence level in the last common ancestor of Brassicaceae (Nelson et al., 2016).

**Figure 1:**
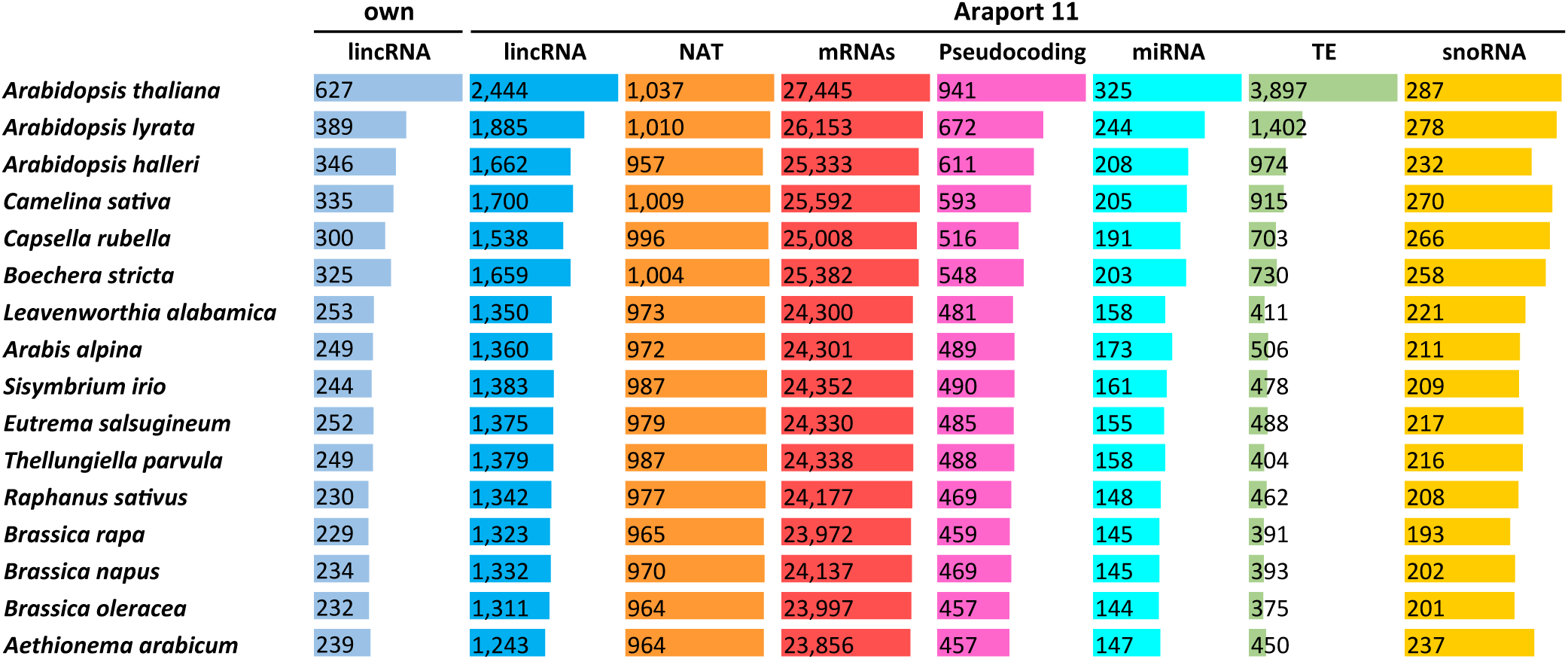
Conservation of genes by position in WGA. *Own:* lincRNAs genes expressed in shade experiments (Kohnen et al., 2016). *Araport11 database annotations (Cheng et al., 2017)*: lincRNAs (long intergenic non-coding RNAs), NAT (Natural antisense transcripts), Coding genes (messenger RNAs), miRNA (microRNAs), Pseudocoding (Pseudocoding genes), TE (Transposable elements), snoRNA (Small nucleolar RNAs)

Conservation of splice sites is a strong indication for the functionality of the transcript. In order to evaluate splice site conservation quantitatively, we constructed a splicing map that identifies for every experimentally determined splice site the homologous position in the other genomes and evaluates them using the MES (see Methods for details). Fig. 2 shows the splicing map for the lincRNA TCONS00053212-00053217 as an illustrative example. Despite the unusually complex transcript structure and the conservation throughout the Brassicaceae, so far nothing is known about the function of this lincRNA. While not all splice sites are represented in all species in the WGA, almost all MES values in this lincRNA are well above the threshold of *MES* > 0. This contrasts with a random sampling of splice sites in coding and non-coding regions in all genomes in the WGA (Suppl. Fig. S4). Indeed, the probability of identifying a random splice site with an MES value greater than 0 in *A. thaliana* is 0.0237 (acceptor) and 0.0165 (donor) for coding genes, and 0.0225 (acceptor) and 0.0168 (donor) for lincRNAs (Suppl. Fig. S4). Most of this lincRNA isoforms therefore can be expected to be present throughout the Brassicaceae, even though the locus is not annotated in Ensembl Plants (release 42) for *B. oleracea, B. rapa*, and *A. lyrata*. Only the short first exon and the 5’ most acceptor of the last exon are poorly conserved by sequence even in close relatives of *A. thaliana*.

**Figure 2:**
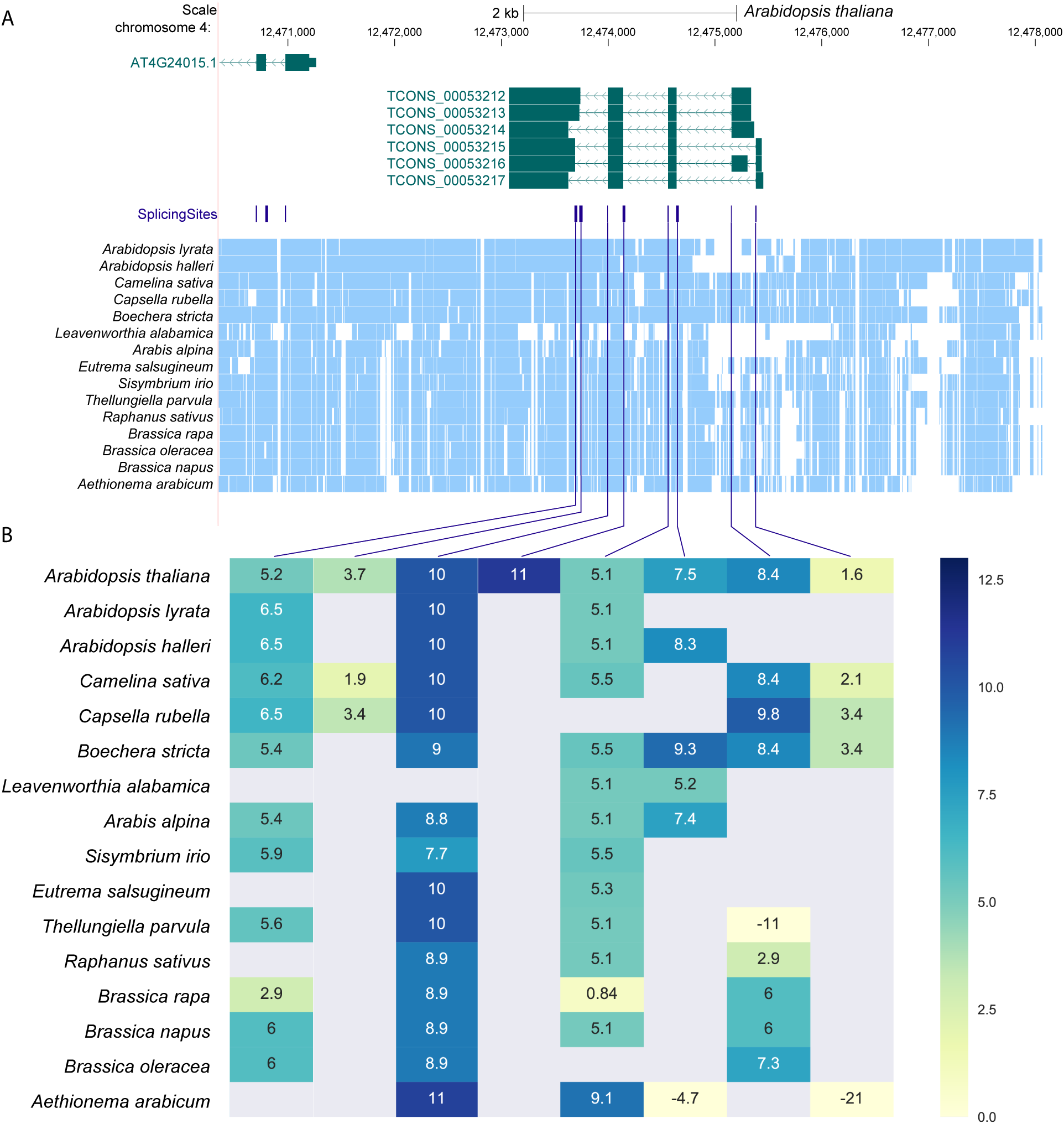
Splicing conservation map of lincRNA locus TCONS00053212-TCONS00053217. **A)** UCSC Genome browser screenshot of the TCONS00053212-TCONS00053217 locus, blocks denote exons and line with arrows introns. The arrow direction indicates direction of transcription. Splicing sites are shown in *purple. Light blue* blocks represent aligned regions as identified by Cactus. **B)** Heatmap of TCONS00053212-TCONS00053217 MES each splice sites (columns) in each species (rows), linked to its position in panel A with a purple line. MES are shown from more negative *(light yellow*) to more positive (*dark blue*). MES values > 0 were used to identify conserved splice sites.

In order to validate the predicted lincRNA splice sites, we investigated publicly available RNA-seq data from eight of the species included in this study (Suppl. Tab. S2). The depth of these data varied considerably. We therefore compared the fraction of recovered lncRNA predictions with the fraction of mRNAs that were detectable in the same RNA-seq data (Suppl. Tab. S4). As expected, we observed that the relative validation rate increases with the depth of data, presumably owing to the fact that lncRNAs are on average less highly expressed and more specifically expressed than mRNAs. Nevertheless, the validation rate in our data of lincRNAs is on average 33.3% in the eight species used for validation and 10.7% for lincRNAs in Araport11 (Cheng et al., 2017), while for coding genes it is 57.5%.

On a genome-wide scale, the conservation of splice sites in lincRNAs provides a lower bound on the fraction of lincRNAs that are under selective constraint as a transcript. We find that 112 of the 173 spliced *A. thaliana* lincRNAs have at least one conserved splice site in another species (Fig. 3).

**Figure 3:**
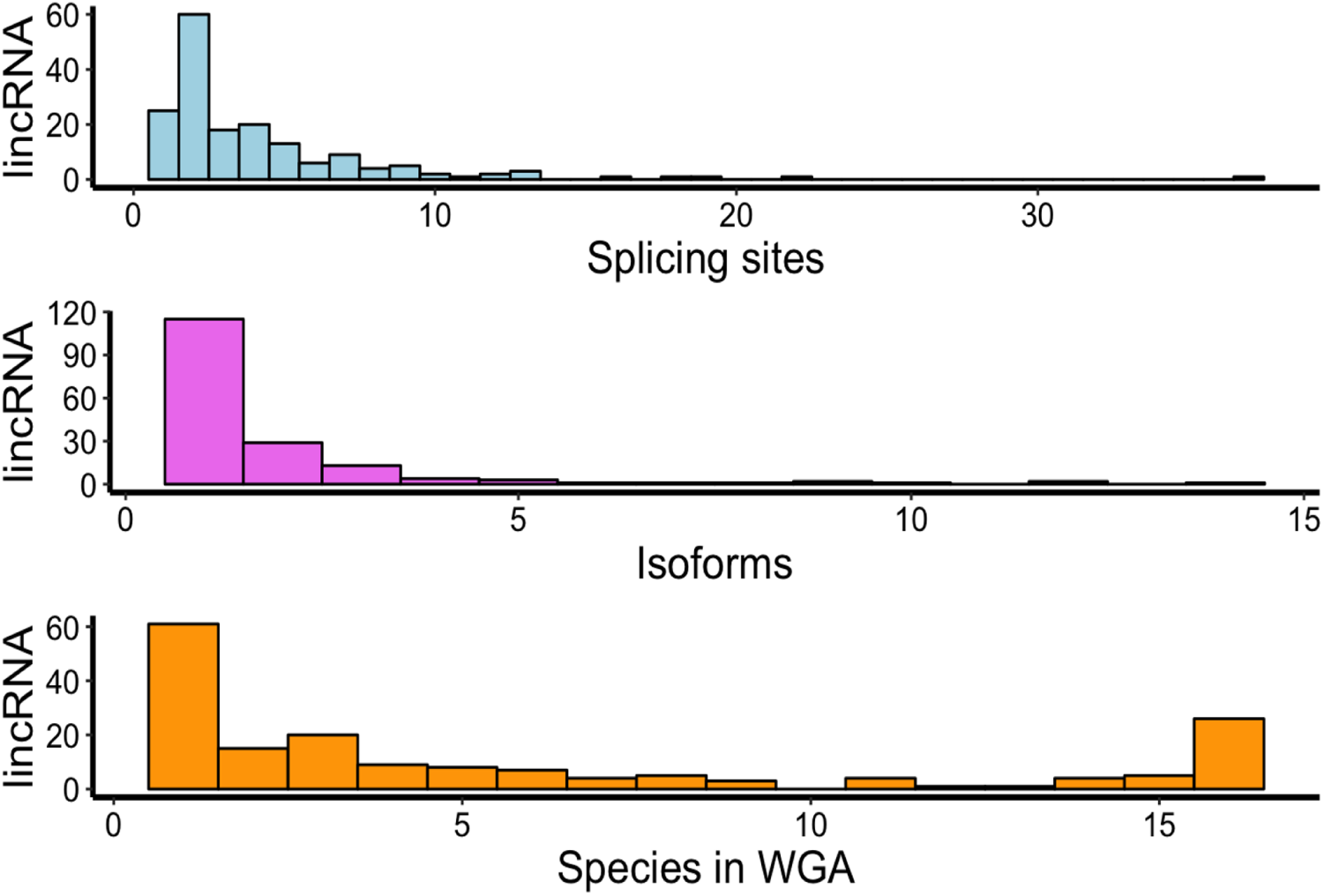
Histograms showing number of splicing sites, isoforms and conservation in WGA of the 173 lincRNAs genes with introns in *Own* dataset. Blue bars indicate the number of splicing sites per lincRNA gene with introns. Purple bars visualize the number of isoforms by lincRNA gene. Orange bars refer to the number of species in which lincRNA gene are conserved.

As expected, we find that splice sites in lincRNAs are much less well conserved than splice sites in protein coding genes (Fig. 4). In total, we identified 39 lincRNAs conserved between the most distant species and *A. thaliana* and 26 lincRNAs with conservation in at least one splice site in the 16 species included in the WGA. These numbers are much lower than for coding genes. Albeit this is expected, given the high conservation of protein coding genes, one has to keep in mind that coding genes on average have at least 6 introns (Deng et al., 2018b), hence it is much more likely to observe conservation of at least one splice site and in lincRNAs with only one or two introns (see Fig. 3).

**Figure 4:**
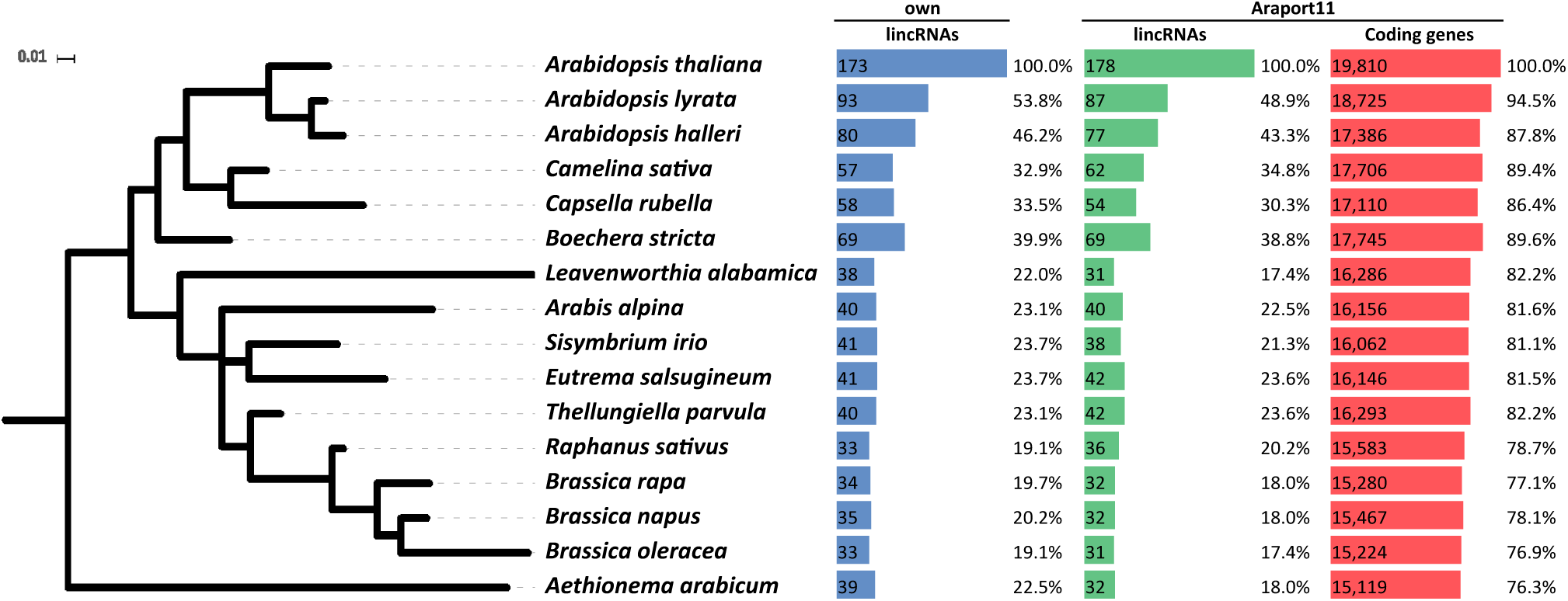
Conservation genes in the Brassicaceae family measured by the conservation of splice sites. *Blue:* own lincRNA set (627); *green:* lincRNAs in Araport11 (2,444); and *red:* coding RNA genes (27,445). Only genes with at least one intron are shown. Phylogenetic tree scale is in changes per site.

The potential incompleteness of annotated lncRNAs, e.g. due to low expression levels, is of concern in this context. It has little influence on our conclusions, however, since incomplete or fragmented annotation only causes us to underestimate the depth of conservation: we might occasionally miss the best-conserved splice junction and we might count fragments as independent, less conserved lncRNAs. Unrecognized overlap with known short ncRNAs is of little concern because the latter are almost never spliced. The only exception are the “splice-site-overlapping” SO-microRNAs (Mattioli et al., 2014), which, however, are almost exclusively found in coding genes (Pianigiani et al., 2018) and thus removed by our filters. We therefore assume that such artefacts have a very minor impact in our analysis.

In comparison to vertebrates, we observe a much lower level of conservation as measured by gene structure. For instance, 35.2% of the transcripts are conserved between human and mouse (Nitsche et al., 2015), while between *A. thaliana* and *A. arabicum* we only find splice site conservation in 6.2% (39/627) of our own lincRNAs and 1.3% (32/2444) of lincRNAs annotated in Araport11. This difference is even more striking given the fact that the evolutionary distance between human and mouse (∼75 Mya) (Waterston et al., 2002) is larger than between *A. thaliana* and *A. arabicum* (∼ 54 Mya) (Beilstein et al., 2010).

Transposable elements (TEs) are important factors in lncRNA origin (Kapusta et al., 2013). To explore if conserved lincRNAs may be related to TEs, we compared our 627 lincRNAs with the genomic positions of TEs described in Araport11 database. We find only 149 of 627 lincRNAs overlap with TEs and these lincRNAs display significantly lower positional conservation than other lincRNAs in the WGA. Indeed, only 11 were found to be positionally conserved between *A. thaliana* and *B. rapa*. The number of TEs which coincide with lincRNAs with conserved splice sites is even smaller; of the 173 lincRNAs with introns only 11 overlapped with TEs. From all 3,897 TEs in the Araport11 database, only 450 are conserved by position in the WGA between *A. thaliana* and *A. arabicum*. This represents only 11.5% of the TEs, i.e., less than the percentage of the lincRNAs conserved by genomic position (Fig. 1).

## 4. DISCUSSION

In this work we explore the conservation of lncRNAs in the Brassicaceae plant family and we find conservation at different levels: from 627 lincRNAs identified we have 38.1% (239/627) conserved by genomic position as determined by the presence of alignable sequence. A small fraction (27.6%) of these lincRNAs contain introns. Only 19.1% of spliced lincRNAs are conserved between *A. thaliana* and *B. oleracea*, the species with the lowest level of conservation in our data set. While sequence conservation may be a consequence of selective constraints on DNA elements, conservation of splice sites directly indicates selective constraints at the transcript level, and thus can be interpreted as evidence for an (unknown) functional role of the lincRNA. The 112 lincRNAs with conserved splice sites are therefore attractive candidates for studies into lncRNA function.

In spite of the small number of spliced lincRNAs analyzed, we find most of them (nearly 65 %) have at least one conserved splice site. This is substantially higher than estimates of conservation by sequence of about ∼22 % amongst Brassica species (Nelson et al., 2016). Thus there is a stronger evolutionary constraint on plant lincRNA processing as measured by splice site conservation than by sequence. This is similar to what was previously found in placental mammals (Nitsche et al., 2015), where about 70% of the lncRNAs have splice site conservation. However, this level of conservation should be considered lower given the divergence time between placental mammals is larger than the divergence times between the Brassicaceae analyzed in this study (52.6 Mya (Kagale et al., 2014)). At least in part this difference is the consequence of the prevalence of single-exon lincRNAs in this clade and the small number of splice sites in those lincRNAs that contain introns. This reduced the power of the method we used to detect splice site conservation, and hints at a reduced importance of introns in the small genomes of the Brassicaceae. The apparent lower conservation of splice sites may also result from our decision to use *A. thaliana* as a reference which, in addition to having a drastically reduced genome, may have also been subjected to clade-specific intron-loss. Transcriptomes of other Brassicas and other plant families that have not undergone drastic genome reduction will help clarify the actual prevalence on monoexonic and intron-gain -loss in plant lncRNAs.

When comparing with other plant families, for example Poaceae, we find that around 20% of maize and rice lincRNAs are conserved by position (Wang et al., 2015), while we find 38.1% (239/627) of lincRNAs conserved in Brassicaceae. These numbers are roughly comparable given that the divergence times of the two families are similar: Brassicaceae, 52.6 Mya (Kagale et al., 2014); Poaceae, 60 Mya (Charles et al., 2009). The lower conservation observed in Poaceae may be explained by the much larger genome size, and thus higher content of repetitive and unconstrained sequences, leaving conserved sequence regions more “concentrated” – and therefore easier to align – in the small genomes of the Brassicaceae. Consistent with previous findings (Nelson et al., 2016), we find that only a small fraction of our lincRNAs associated with TEs, compared to a much stronger association in Poaceae (Wang et al., 2017). We interpret this to be consequence of the substantial reduction of genome size in Brassicas. More detailed comparisons of lincRNA conservation with other plant families will have to await better assembled and annotated genomes to construct adequate WGAs.

A limitation of our work is the restriction to intergenic lncRNAs, caused by the need to avoid potential overlaps of the splice sites with other constrained elements. High quality transcriptomes from diverse tissues for most species could alleviate this shortcoming, allowing us to construct splicing maps using only experimental evidence. Spurious sequence conservation would then no longer influence the results. This is of particular relevance in Brassicaceae, since about 70% of transcripts have antisense lncRNAs (Wang et al., 2014). These had to be excluded from our the analysis even though at least some of them, e.g. *COOLAIR* (Hawkes et al., 2016), are known to have important biological functions. Complementarily to the analysis of splice site conservation, conserved RNA secondary structure can serve as evidence of selection constraints at the RNA level (Washietl et al., 2005). Moreover, structural analysis can be applied to both spliced and monoexonic transcripts. So far, no genome-wide assessment of conservation of RNA secondary structure has been reported for plants. However, recent structurome sequence data indicates that RNA structure is also under selection at the genome-wide level in plants (Deng et al., 2018a).

In summary we showed here that higher plants contain at least dozens and most likely hundreds of wellconserved – and with near certainty functional – long non-coding RNAs. We provide an initial catalog of candidates for more detailed exploration, in many cases supported by direct evidence for expression in several species. We furthermore contribute a generic workflow that can be used to uncover conserved lncRNAs in other groups of plants. Given the rapidly expanding collection of publicly available RNA-seq data sets, we suggest that a comparative analysis of lncRNA conservation can complement standard procedures for genome annotation and thus eventually lead to comprehensive picture of lncRNA diversity and evolution in plants.

## ACKNOWLEDGMENTS

This work was funded in part by CONACYT (JAC-G: CONACYT Ph.D. Scholarship - 338379, SLF-V: CONACYT Research Fellowship 2015-72223), by the *Deutsche Forschungsgemeinschaft* (DFG): grant no. STA850/19-1 to PFS and by a Royal Society Newton Advanced Fellowship (NAF\R1\180303) awarded to SLF-V. We are grateful to Thomas Gatter for advice on transcriptome assembly.

