## Supplemental Material PDF for "Splicing conservation signals in plant long non-coding RNAs"

Table S1: **Genome versions used in this study.**

| Species | Ensembl ID | Source |
|---|---|---|
| *Aethionema arabicum* | GCA_000411095.1 | NCBI |
| *Arabidopsis halleri* | GCA_900078215.1 | Phytosome v12 |
| *Arabidopsis lyrata* | GCA_000004255.1 | Ensembl-Plants |
| *Arabidopsis thaliana* | GCA_000001735.1 TAIR10 | Ensembl-Plants |
| *Arabis alpina* | GCA_000733195.1 | NCBI |
| *Boechera stricta* | GCA_002079875.1 | NCBI |
| *Brassica napus* | GCA_000686985.1 | Ensembl-Plants |
| *Brassica rapa* | GCA_000309985.1 | Ensembl-Plants |
| *Camelina sativa* | GCA_000633955.1 | NCBI |
| *Capsella rubella* | GCA_000375325.1 | Phytosome v12 |
| *Leavenworthia alabamica* | GCA_000411055.1 | NCBI |
| *Raphanus sativus* | GCA_000801105.2 | NCBI |
| *Thellungiella parvula* | GCA_000218505.1 | NCBI |
| *Sisymbrium irio* | GCA_000411075.1 | NCBI |
| *Brassica oleracea* | GCA_000695525.1 | Ensembl-Plants |
| *Eutrema salsugineum* | GCA_000478725.1 | NCBI |

*Corresponding authors

Table S2: **Transcriptomes used in this study**

| Organism | Organism part | RUN | Link |
|---|---|---|---|
| *Camelina sativa* | Whole Shoot | SRR5920412 | www.ncbi.nlm.nih.gov/geo/query/acc.cgi?acc=GSM2736329 |
| *Boechera stricta* | Whole plant | SRR3178610 | www.ncbi.nlm.nih.gov/geo/query/acc.cgi?acc=GSM2067369 |
| *Arabis alpina* | Leaf | SRR5004109 | www.ncbi.nlm.nih.gov/geo/query/acc.cgi?acc=GSM2385855 |
| *Raphanus sativus* | Whole shoot | SRR8506403 | www.ncbi.nlm.nih.gov/geo/query/acc.cgi?acc=GSM3583822 |
| *Brassica rapa* | Leaf | SRR2060322 | www.ncbi.nlm.nih.gov/geo/query/acc.cgi?acc=GSM1708760 |
| *Eutrema salsugineum* | Leaf | SRR2922646, SRR2922647, SRR2922648, SRR2922649 | www.ncbi.nlm.nih.gov/geo/query/acc.cgi?acc=GSM1942123 |
| *Aethionema arabicum* | Seed | SRR8503241 | www.ncbi.nlm.nih.gov/sra/SRX5307165[accn] |
| *Arabidopsis thaliana* | Roots, leaves, flowers and siliques | SRR505743, SRR505744, SRR505745, SRR505746 | www.ebi.ac.uk/gxa/experiments/E-GEOD-38612/ |
| *Brassica oleracea* | Roots, leaves, flowers, callus, fruit, steam, flower bud | SRR630922, SRR630923, SRR630924, SRR630925, SRR630926, SRR630927, SRR630928 | www.ebi.ac.uk/gxa/experiments/E-GEOD-42891/Results |

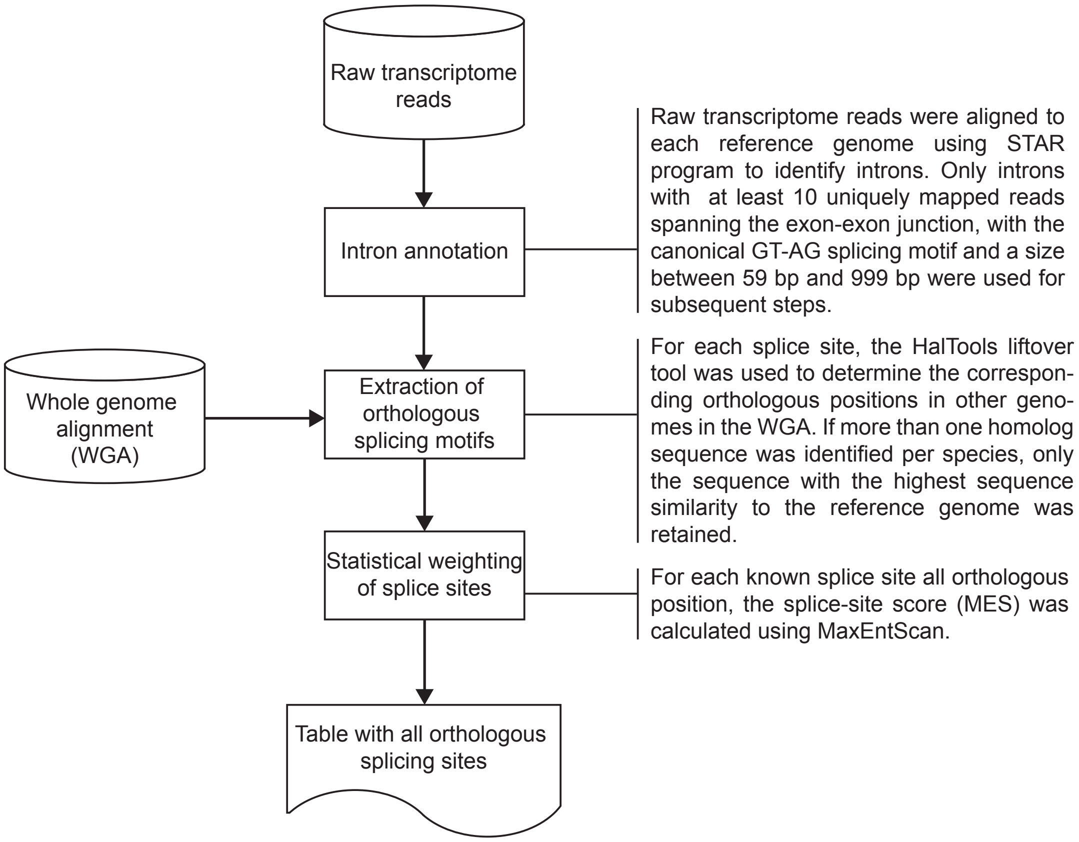


Figure S1: **Splice map bioinformatics pipeline** Flow chart displaying the three main steps to generate a splice map. Full pipeline available at: bitbucket.org/JoseAntonioCorona/splicing_map_plants.

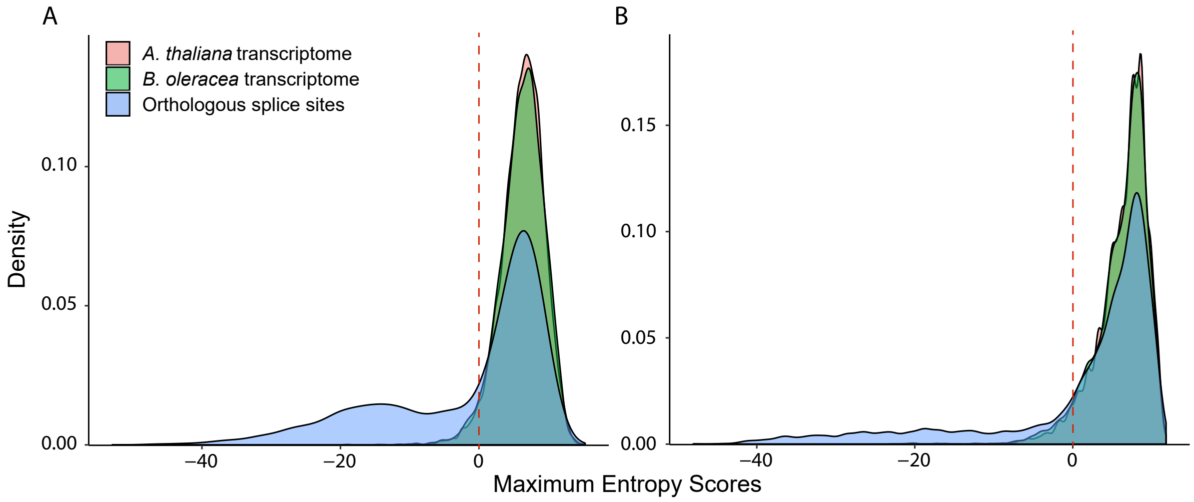


Figure S2: **Density distribution of MES** for splice sites in the donor (A) and acceptor (B) site as identified in *A. thaliana* (*pink*) and *B. oleracea* (*green*) transcriptomes compared to orthologous splice sites (*blue*) identified by position in the Cactus generated WGA. The red line denotes the Maximum Entropy threshold used to consider a splice site as real in the alignment (MES > 0).

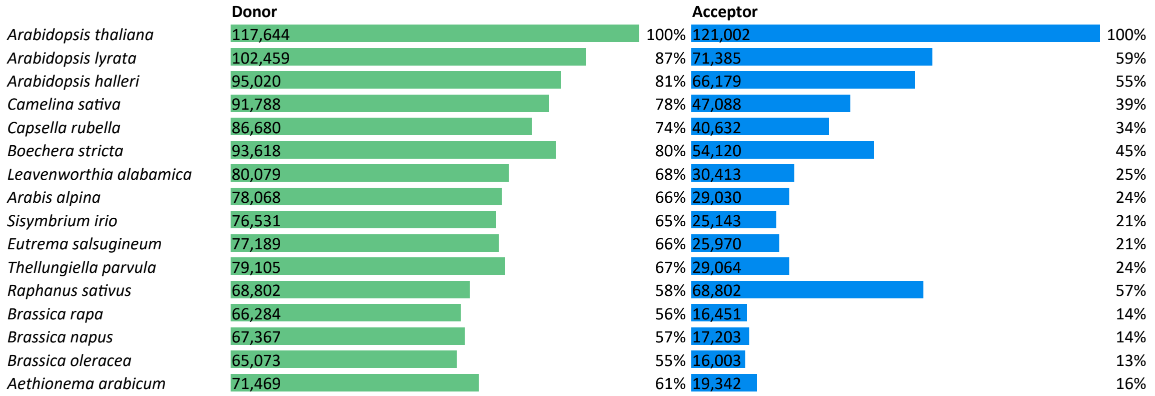


Figure S3: **Conservation of splice sites** Number of *A. thaliana* splice sites identified as conserved in the rest of the species according to the WGA.

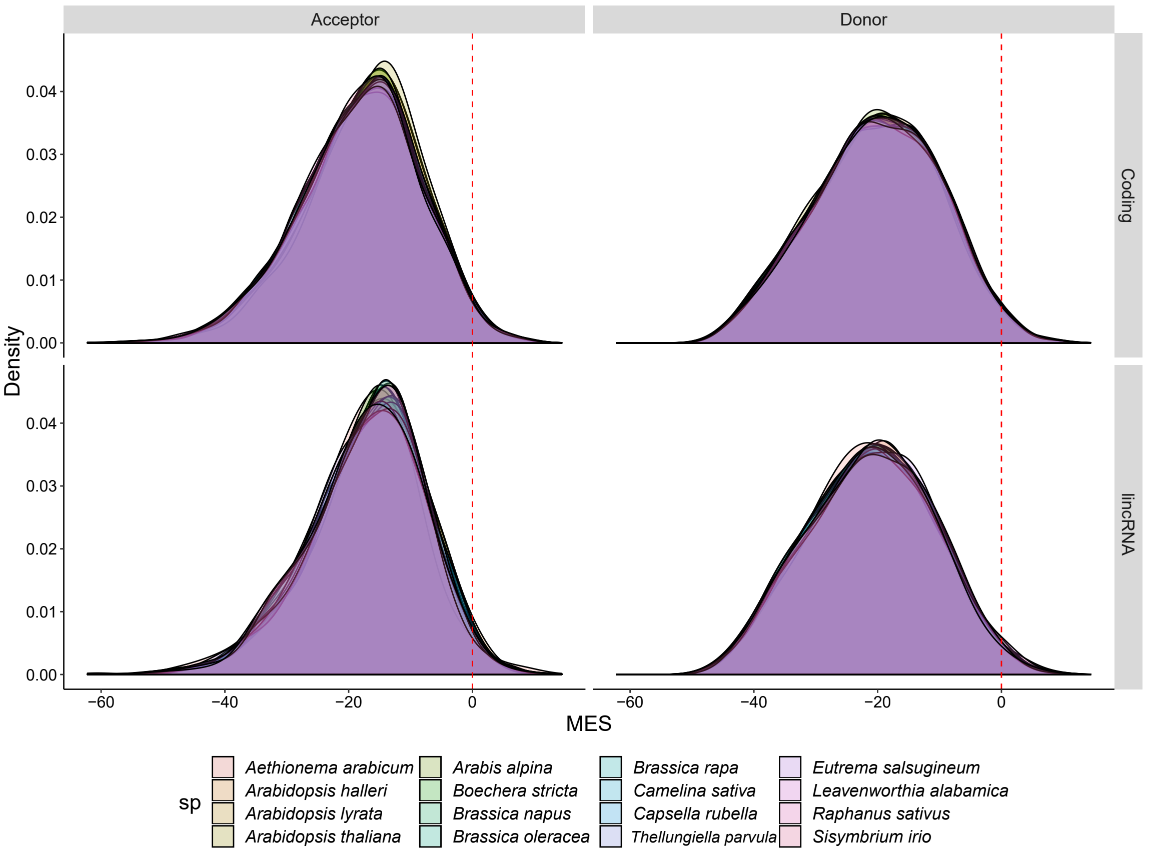


Figure S4: **Density distribution MES for random positions** Density distribution of the probability of finding random splice sites along coding genes (top panels) and lincRNAs (bottom panels). We used 10,000 random splice positions for both acceptor (left panels) and donor (right panels) motifs for all species in the WGA.

Table S3: **Annotation of splicing sites in different species** Expression at the splicing sites using the available transcriptomes of the WGA species listed in Table S2. The number of splicing sites preserved by position with respect to *A. thaliana* are shown in the yellow bars.

| | Splicing sites | | |
|---|---|---|---|
| | **Total** | **Conserved in WGA** | |
| ***Arabidopsis thaliana*** | 222,772 | 222,772 | 100% |
| ***Camelina sativa*** | 64,672 | 35,925 | 16% |
| ***Boechera stricta*** | 103,337 | 95,935 | 43% |
| ***Arabis alpina*** | 72,424 | 62,883 | 28% |
| ***Eutrema salsugineum*** | 36,121 | 32,073 | 14% |
| ***Raphanus sativus*** | 36,179 | 27,893 | 13% |
| ***Brassica rapa*** | 11,664 | 8,843 | 4% |
| ***Brassica oleracea*** | 97,674 | 61,843 | 28% |
| ***Aethionema arabicum*** | 26,894 | 22,665 | 10% |

Table S4: **Splice-site validation in transcriptomes in other species** Genes with splicing are shown (lincRNAs: own set (173); Araport11 (178); coding genes: Araport11 (19,810)). Genes whose splice sites are expressed and conserved both in *A. thaliana* and in the corresponding species are listed in the *validated* column. The *lincRNA/mRNA* column denotes the percentage of validated lincRNAs over the percentage of validated coding mRNAs.

| Organism | own | | | | Araport11 | | | | | | |
|---|---|---|---|---|---|---|---|---|---|---|---|
| | lincRNAs *A. thaliana* | validated | | lincRNA/mRNA | lincRNAs *A. thaliana* | validated | | lincRNA/mRNA | mRNAs *A. thaliana* | validated | |
| ***Arabidopsis thaliana*** | 173 | 173 | 100.0% | 100.0% | 178 | 178 | 100.0% | 100.0% | 19,810 | 19,810 | 100.0% |
| ***Camelina sativa*** | 57 | 15 | 26.3% | 55.6% | 62 | 5 | 8.1% | 17.0% | 17,706 | 8,383 | 47.3% |
| ***Boechera stricta*** | 69 | 27 | 39.1% | 55.0% | 69 | 18 | 26.1% | 36.7% | 17,745 | 12,624 | 71.1% |
| ***Arabis alpina*** | 40 | 17 | 42.5% | 59.8% | 40 | 7 | 17.5% | 24.6% | 16,156 | 11,490 | 71.1% |
| ***Eutrema salsugineum*** | 41 | 7 | 17.1% | 31.2% | 42 | 4 | 9.5% | 17.4% | 16,146 | 8,832 | 54.7% |
| ***Raphanus sativus*** | 33 | 10 | 30.3% | 60.0% | 36 | 2 | 5.6% | 11.0% | 15,583 | 7,865 | 50.5% |
| ***Brassica rapa*** | 34 | 7 | 20.6% | 82.4% | 32 | 3 | 9.4% | 37.5% | 15,280 | 3,818 | 25.0% |
| ***Brassica oleracea*** | 33 | 23 | 69.7% | 78.0% | 31 | 0 | 0.0% | 0.0% | 15,224 | 13,598 | 89.3% |
| ***Aethionema arabicum*** | 39 | 8 | 20.5% | 40.3% | 32 | 3 | 9.4% | 18.4% | 15,119 | 7,693 | 50.9% |
| **Average (excluding *A. thaliana* )** | | 14.25 | 33.3% | | | 5.25 | 10.7% | | | 9287.9 | 57.5% |

**Dataset 1:**
TrackHubs of WGA, including annotation of splicing sites, Araport11 annotations and own lincRNA annotation.
www.bioinf.uni-leipzig.de/Publications/SUPPLEMENTS/19-001/BrassicaceaeWGA/hub.txt

**Dataset 2:**
Table of the splice sites, the table contains all the splicing sites that we have predicted for *A. thaliana* and their homologs by position in the 15 species of the WGA. http://www.bioinf.uni-leipzig.de/Publications/SUPPLEMENTS/19-001/SplicingMap.tsv

**Dataset 3:**
Scripts used in creation of splicing map.
bitbucket.org/JoseAntonioCorona/splicing_map_plants.

**Dataset 4:**
Conservation table by position of TE and lincRNAs overlapping TE.
http://www.bioinf.uni-leipzig.de/Publications/SUPPLEMENTS/19-001/lincRNA-overlap-TE.tsv

**Dataset 5:**
BED files of our lincRNAs for *A. thaliana* and their homologs by genomic position in the 15 species of the WGA
http://www.bioinf.uni-leipzig.de/Publications/SUPPLEMENTS/19-001/lincRNAs-position/

**Dataset 6:**
LincRNAs with evidence of expression in transcriptomes of different species.
https://www.bioinf.uni-leipzig.de/Publications/SUPPLEMENTS/19-001/lincRNA_Araport_expression_by_specie.tsv
https://www.bioinf.uni-leipzig.de/Publications/SUPPLEMENTS/19-001/lincRNA_own_expression_by_specie.tsv
